# Uncoupling sodium channel dimers rescues phenotype of pain-linked Nav1.7 mutation

**DOI:** 10.1101/716654

**Authors:** Annika H. Rühlmann, Jannis Körner, Nikolay Bebrivenski, Silvia Detro-Dassen, Petra Hautvast, Carène A. Benasolo, Jannis Meents, Jan-Philipp Machtens, Günther Schmalzing, Angelika Lampert

## Abstract

The voltage-gated sodium channel Nav1.7 is essential for an adequate perception of painful stimuli. Its mutations cause various pain syndromes in human patients. The hNav1.7/A1632E mutation induces symptoms of erythromelalgia and paroxysmal extreme pain disorder (PEPD), and its main gating change is a strongly enhanced persistent current.

Using molecular simulations, we demonstrate that the disease causing persistent current of hNav1.7/A1632E is due to impaired binding of the IFM motif, thus affecting proper function of the recently proposed allosteric fast inactivation mechanism. By using native polyacrylamide gel electrophoresis (PAGE) gels, we show that hNav1.7 dimerizes. The disease-linked persistent current depends on the channel’s functional dimerization status: Using difopein, a 14-3-3 inhibitor known to uncouple dimerization of hNav1.5, we detect a significant decrease in hNav1.7/A1632E induced persistent currents.

Our work identifies that functional uncoupling of hNav1.7/A1632E dimers rescues the pain-causing molecular phenotype by interferes with an allosteric fast inactivation mechanism, which we link for the first time to channel dimerization. Our work supports the concept of sodium channel dimerization and reveals its relevance to human pain syndromes.

## Introduction

Voltage-gated sodium channels (VGSC/Nav) play a crucial role in the perception and transduction of painful signals (Ahern *et al*, 2016) and mutations leading to chronic pain syndromes severely affect the patients’ quality of life (Lampert *et al*, 2010). Human VGSC consist of four domains (DI-DIV), each containing six transmembrane segments (S1-S6) (Shen *et al*, 2017; Yan *et al*, 2017; Pan *et al*, 2018; Clairfeuille *et al*, 2019; Xu *et al*, 2019; Shen *et al*, 2019). The ion pore is formed by the S5 and S6 segments of each domain, and it is opened by tethered voltage-sensing parts, formed by S1–S4, which move outward upon depolarization (Catterall, 2014; Guy & Seetharamulu, 1986; Sula *et al*, 2017; Payandeh *et al*, 2011). Within milliseconds after pore opening the VGSCs inactivate: Three amino acids located in the DIII–DIV linker were identified to form the so-called IFM motif or inactivation particle.

Until very recently, it was thought that the IFM motif blocks ion permeation by binding to the cytoplasmic side of the pore, thus causing fast inactivation (Armstrong *et al*, 1973; West *et al*, 1992). This process has been described as “hinged lid” mechanism (Eaholtz *et al*, 1994; West *et al*, 1992). However, with the recently published high-resolution structure of VGSC subtypes NavPas, Nav1.4 and Nav1.7, a new fast inactivation mechanism has been proposed: Instead of a direct occlusion of the pore by the IFM motif, the inactivation particle may bind to a hydrophobic binding pocket in the periphery of the S6 helical bundle and thus cause fast inactivation due to an allosteric effect. This induces movement of the DIVS6 helix towards the ion permeation pathway. At the same time DIIIS6 is pulled towards the ion pore, possibly causing the DIS6 and DIIS6 helices to move in the same direction to close the inner gate of the permeation pathway (Pan *et al*, 2018; Yan *et al*, 2017; Xu *et al*, 2019; Clairfeuille *et al*, 2019; Shen *et al*, 2019, 2017).

In humans, nine different VGSC isoforms are identified (Ahern *et al*, 2016). Recent studies suggest that the cardiac subtype Nav1.5 forms functionally coupled dimers (Clatot *et al*, 2017, 2018). This dimerization seems to be mediated by 14-3-3, an abundantly expressed protein also involved in enzyme regulation and neurological diseases (Foote & Zhou, 2012; Obsil *et al*, 2001; Clatot *et al*, 2017). 14-3-3 presents two binding grooves which bind to specific motifs, often marked by a sequence containing phosphoserine or other phosphorylated amino acids (Obsil & Obsilova, 2011; Johnson *et al*, 2010). For hNav1.5, it was suggested that 14-3-3 binds in the linker region between DI and DII (Clatot *et al*, 2017). There are seven different 14-3-3 isoforms, all of which can be inhibited by the artificial protein difopein (*di*meric-*fo*urteen-three-three-*pe*ptide *in*hibitor)(Masters & Fu, 2001). Difopein blocks 14-3-3 and is therefore suggested to be able to inhibit functional coupling of VGSC dimers (Clatot *et al*, 2017; Masters & Fu, 2001).

In the present study, we focus on the pain-related VGSC subtype Nav1.7, which is expressed in nociceptive sensory neurons and free nerve terminals in the skin (Djouhri *et al*, 2003; Black *et al*, 2012; Persson *et al*, 2010). Mutations of Nav1.7 are linked to pain syndromes, such as inherited erythromelalgia (IEM) or paroxysmal extreme pain disorder (PEPD) (Lampert *et al*, 2010). Additionally, patients with complete loss-of-function mutations of Nav1.7 show strongly impaired pain perception (congenital insensitivity to pain, CIP) (Cox *et al*, 2006; McDermott *et al*, 2019), stressing the important role of Nav1.7 in the generation of human pain.

We study the previously reported gain-of-function mutation hNav1.7/A1632E mutation, which induces a combination of IEM and PEPD in the heterozygous carrier and is characterized by an incomplete fast inactivation leading to a prominent persistent current (Estacion *et al*, 2008; Eberhardt *et al*, 2014). Several natural occurring and artificially induced mutations at this position were reported, showing that this area of the VGSC protein is involved in regulation of voltage dependence and kinetics of fast inactivation (Eberhardt *et al*, 2014; Yang *et al*, 2016). Using molecular dynamics simulations, we demonstrate that the increase of size and addition of negative charge at position 1632 impedes binding of the IFM motif to a hydrophobic binding pocket, thus preventing fast inactivation, supporting the new allosteric inactivation mechanism suggested by the recent cryo-EM structures (Pan *et al*, 2018; Yan *et al*, 2017; Shen *et al*, 2017, 2019; Clairfeuille *et al*, 2019). Using the CIP-mutations hNav1.7/R896Q and hNav1.7/G375Afs (Cox *et al*, 2010; Shorer *et al*, 2014), we show that hNav1.7 dimerizes and we find evidence that this channel-dimerization affects the allosteric fast inactivation mechanism also in unmutated WT channels. In addition, we present data that reveal how functionally uncoupling dimerization significantly reduces the size of the disease-relevant persistent current of hNav1.7/A1632E.

## Results

### hNav1.7/A1632E impairs fast inactivation – evidence for an allosteric mechanism

Using patch-clamp of HEK cells overexpressing the pain-linked mutation hNav1.7/A1632E, we confirmed its reported prominent persistent current (Fig. 1A-C), suggesting an impaired fast inactivation as major disease-linked gating change (Dib-Hajj *et al*, 2013).

**Figure 1:**
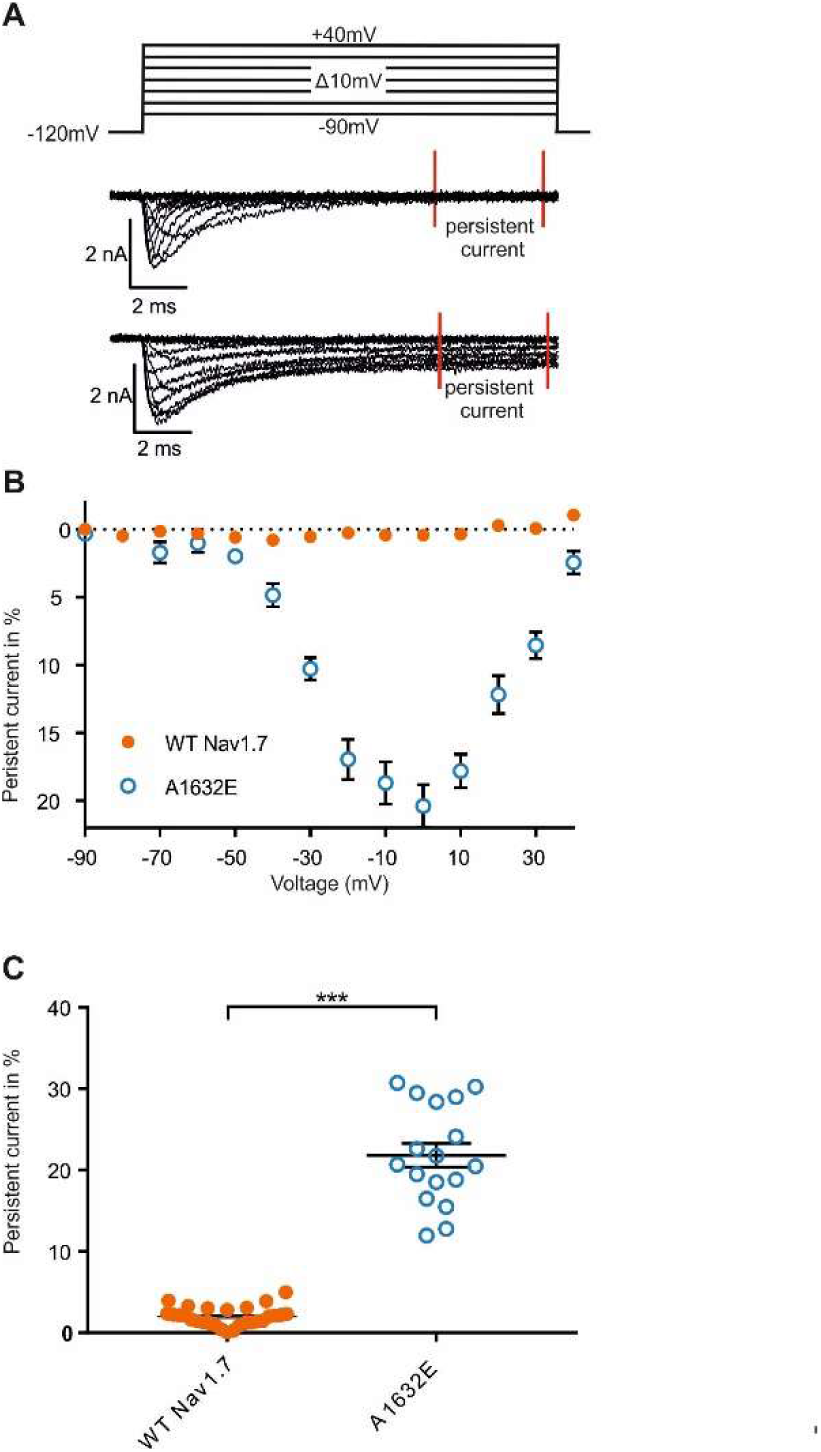
hNav1.7/A1632E presents a prominent persistent current. A) Representative traces of WT and A1632E sodium current. Persistent current was measured in the indicated range (red bars). Voltage-protocol depicted above. B) Relative persistent current in percent of peak current for each test-pulse voltage. The measured persistent current at each voltage step was normalized towards the peak inward current of the cell. C) Maximal relative persistent current of WT and hNav1.7/A1632E. The maximal persistent current of each cell is shown in this panel. Max persistent current: WT: 1.99%±0.22%, N=28; hNav1.7/A1632E: 21.82%±1.47%, N=17. ***p<0,0001. Data are shown as mean ± SEM.

The recently published cryo-EM structures of human Nav1.4 and Nav1.7 revealed the IFM motif for fast inactivation to reside in-between the S6 helical constriction of the inner pore domain and VSDIII (Shen *et al*, 2019; Pan *et al*, 2018). The inactivation particle is tightly bound to a hydrophobic cavity, which is formed by residues from the DIIIS4–S5 linker, DIIIS5, DIIIS6, DIVS5, and DIVS6. A1632 on the DIVS5 is positioned in close proximity (Fig. 2A), and substitution of A1632 by glutamate may affect binding of the IFM motif. To test this hypothesis, we modelled the A1632E substitution in the Nav1.7 structure. The large glutamate side chain protrudes into the binding pocket and could therefore cause steric clashes with the IFM motif (Fig. 2B). However, since substitution of A1632 with D, which is smaller than E, has been shown to induce even larger persistent currents than A1632E (Eberhardt *et al*, 2014), steric repulsion of the IFM motif is unlikely to be the only explanation for these mutants’ effects.

**Figure 2:**
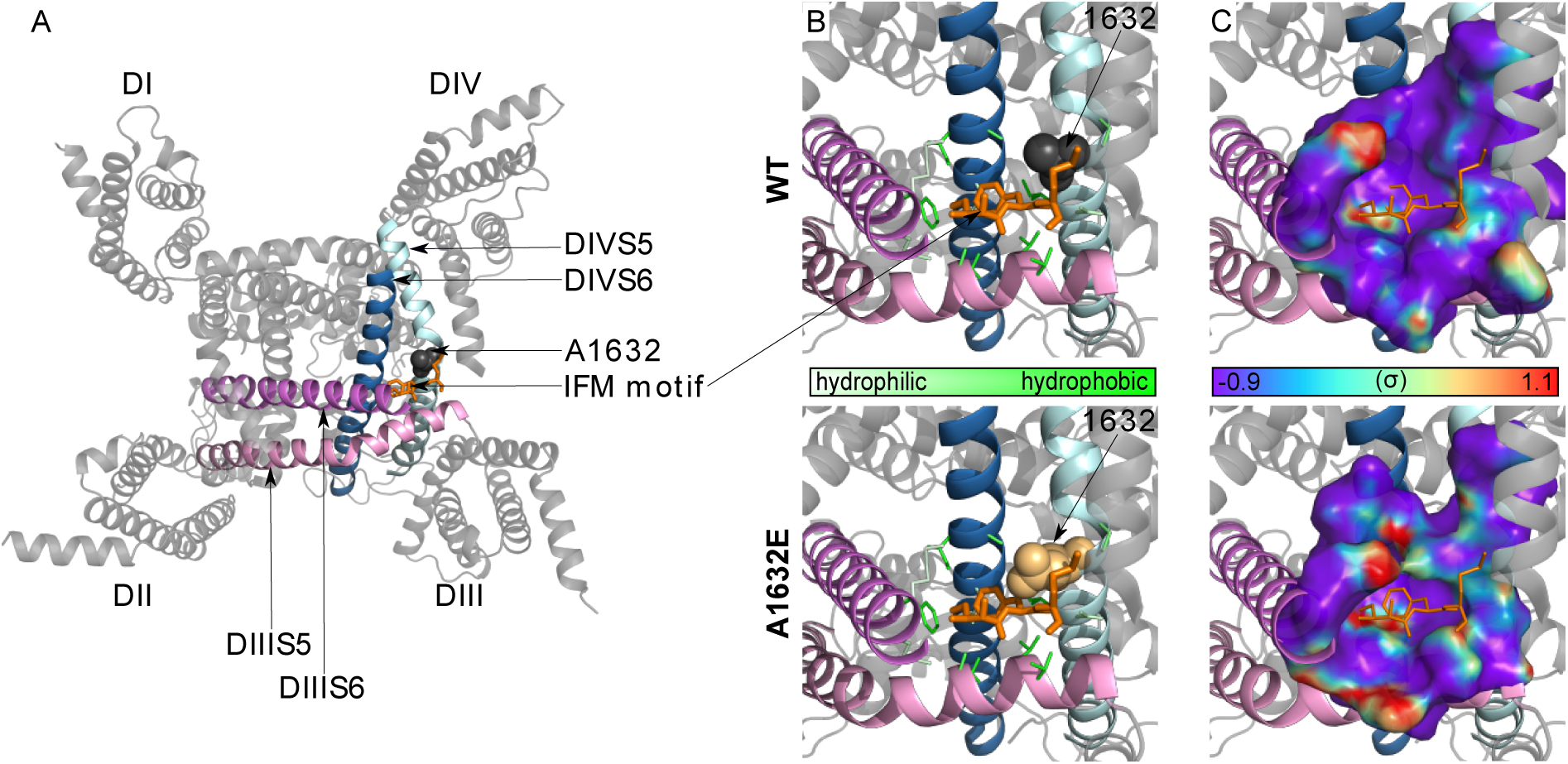
The A1632E mutation impedes binding of the IFM motif via steric repulsion and hydrophobic mismatch. A) Overview of the cryo-EM structure of the hNav1.7 alpha subunit from the intracellular side (PDB ID: 6J8I). A1632 of DIVS5 is shown as spheres, the IFM motif is shown as orange sticks. B) Close-up on the IFM-binding pocket, a hydrophobic cavity formed by residues from the DIIIS4–S5 linker, DIIIS5, DIIIS6, DIVS5, and DIVS6 in WT and A1632E Nav1.7 (thin sticks, pocket-forming residues; thick sticks, IFM-motif residues; spheres, residue 1632). C) Solvent-accessible surface of the IFM-binding pocket from the same perspective as in (B). Water densities from unguided 200-ns all-atom MD simulations of WT and A1632E Nav1.7, with the DIII–DIV linker being removed, are mapped onto the protein surface as defined by the color-scale bar. The IFM motif was thus not present in these simulations and its position in the IFM-bound structure is indicated by orange lines.

Due to the chemical nature of the IFM motif, tight binding of these residues requires a hydrophobic binding pocket, and this property could be disrupted by increased attraction of water molecules into the pocket caused by the hydrophilic A1632E substitution. We performed unguided all-atom molecular dynamics simulations to measure the water accessibility of the empty IFM-binding pocket. Whereas the hydrophobic nature of most sidechains lining the WT binding pocket results in a rather low water accessibility, the A1632E mutation resulted in an increase in hydration at the IFM-binding position (Fig. 2C). Thus, both steric repulsion and hydrophobic mismatch of the binding pocket with the IFM motif contribute to the functional phenotype of A1632E by preventing binding of the IFM motif. Taking into account the recently proposed allosteric inactivation mechanism (Yan et al, 2017), this constellation predicts impaired fast inactivation, resulting in persistent current for the hNav1.7/A1632E mutation: the larger glutamic acid and the increased hydration of the binding pocket impairs the binding of the IFM motif. Thus, in the mutant channel, DIVS6 may be less prone to move into the ion permeation pathway and the pore may therefore not inactivate properly.

### Voltage-gated sodium channel subtype Nav1.7 dimerizes

Recently, it was reported that hNav1.5 forms functional channel-dimers mediated by 14-3-3, which is also abundantly expressed in HEK cells (Supplementary Figure S1 and (Clatot *et al*, 2017, 2018)). Difopein interferes with 14-3-3 binding and can hence be used to inhibit functional coupling of the channel dimers (Clatot *et al*, 2017). To test if oligomerization also occurs in hNav1.7, we solubilized hNav1.7^GFP^-expressing oocytes in the non-ionic detergent digitonin and resolved the protein by SDS-urea-PAGE and hrCN-PAGE. As a possible positive control we co-analyzed hNav1.5^GFP^. Both hNav1.5^GFP^ and hNav1.7^GFP^ migrated in the SDS-PAGE gel at 250 kDa and 220 kDa, in reasonable agreement with the sequence-calculated masses of 256 and 254 kDa, respectively, both in the non-reduced and the reduced states (Fig. 3A). These results argue against the existence of disulfide-bonded VGSC dimers, but non-covalently bound dimers can still exist.

**Figure 3:**
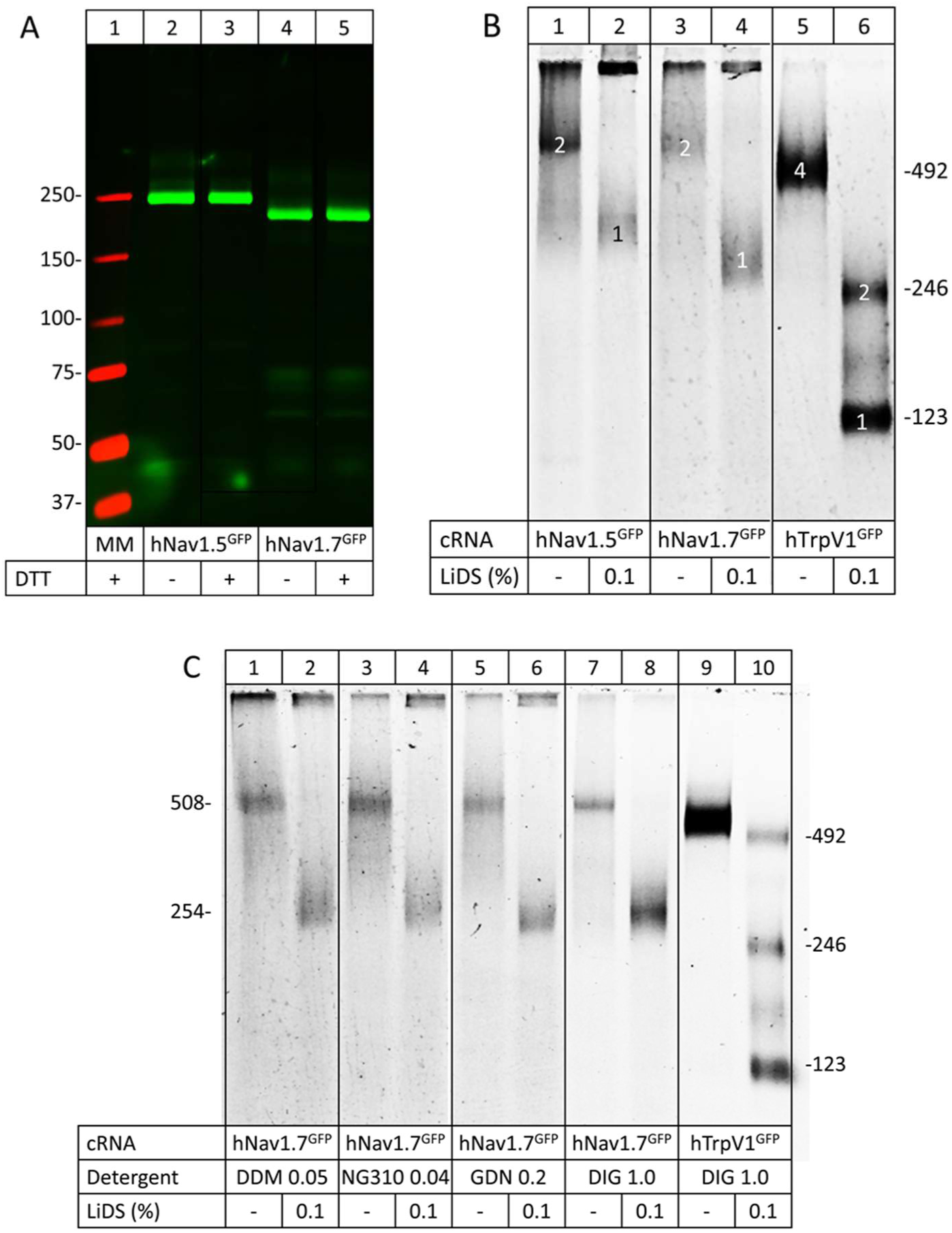
Nav1.5 and Nav1.7 migrate as non-covalent homodimers in native PAGE gels. A) SDS-urea-PAGE of hNav1.5 and hNav1.7. B) and C) hrCN-PAGE of hNav1.5 and hNav1.7 using various detergents to dissolve dimers into monomers. The indicated channel proteins were extracted with digitonin (A, B) or one of the indicated detergents (C) from X. laevis oocytes, resolved by hrCN-PAGE and visualized via their GFP fluorescence by Typhoon fluorescence scanning. Protein migration is shown both under native conditions and after partial denaturation following a 1-h incubation with 0.1% LiDS at 37 °C, as indicated. The numbers in the right margins in (B) and (C) indicated the sequence-calculated masses (protomers to tetramers) of the partially denatured and accordingly partially disassembled hTrpV1GFP channel. The numbers in the left margin in (C) correspond to the sequence-calculated masses of the hNav1.7GFP protomer and homodimer. The abbreviations used are DDM, NG310, GDN, and DIG for dodecylmaltoside, lauryl maltose neopentyl glycol, glyco-diosgenin, and digitonin, respectively.

When the same samples were resolved by high resolution Clear Native-PAGE (hrCN-PAGE), both hNav1.5^GFP^ and hNav1.7^GFP^ migrated under non-denaturing conditions at masses of about 560 kDa (Fig. 3B, lanes 1 and 3). We compared our results with the migration of the co-resolved hTrpV1^GFP^, a 492 kDa homotetramer (Julius, 2013) consisting of four protomers with a calculated mass of 123 kDa each (Fig. 3B). Treatment with a low concentration of 0.1 % of the denaturing detergent lithium dodecyl sulfate (LiDS) resulted in a dissociation of the hTrpV1^GFP^ homotetramer into the homodimer and the protomer with calculated masses of 246 and 123 kDa, respectively (Fig. 3B, lanes 5 and 6). Also, the VGSC proteins dissociated into faster migrating species. Based on the migration of the mass marker hTrpV1^GFP^, hNav1.7^GFP^ migrated at ∼570 kDa and ∼290 kDa in the absence and presence of LiDS, respectively (Fig. 3B, lanes 3 and 4). We conclude from these results that both hNav1.5 and hNav1.7 have a strong propensity to assemble in *X. laevis* oocytes as homodimers.

To address a contribution of the nature of the solubilization detergent on the oligomeric state, we solubilized hNav1.7^GFP^-expressing oocytes in three additional non-ionic detergents, NG310, GDN and digitonin besides DDM. hrCN-PAGE resolved identical migration patterns, consistent with homodimeric and monomeric states in the absence or presence of the denaturing detergent LiDS, respectively (Fig. 3C). Thus, our data clearly indicate that apart from hNav1.5, also hNav1.7 is able to form homodimers.

To verify channel dimerization on a functional level in hNav1.7, we used two loss-of-function mutations reported to occur in CIP patients: hNav1.7/R896Q and hNav1.7/G375Afs (Fig. 4A and (Cox *et al*., 2010; Shorer *et al*., 2014)). The latter causes a truncation of the channel and resides before the proposed dimerization site (Fig. 4A and (Clatot *et al*, 2017)), thus suggesting that these mutant channels would not be able to interact with WT Nav1.7. hNav1.7/R896Q, on the other hand, is supposed to form complete channels, and should thus also interact as a dimer with Nav1.7 WT (see Supplementary Figure S2 for an alignment of the putative dimerization site).

**Figure 4:**
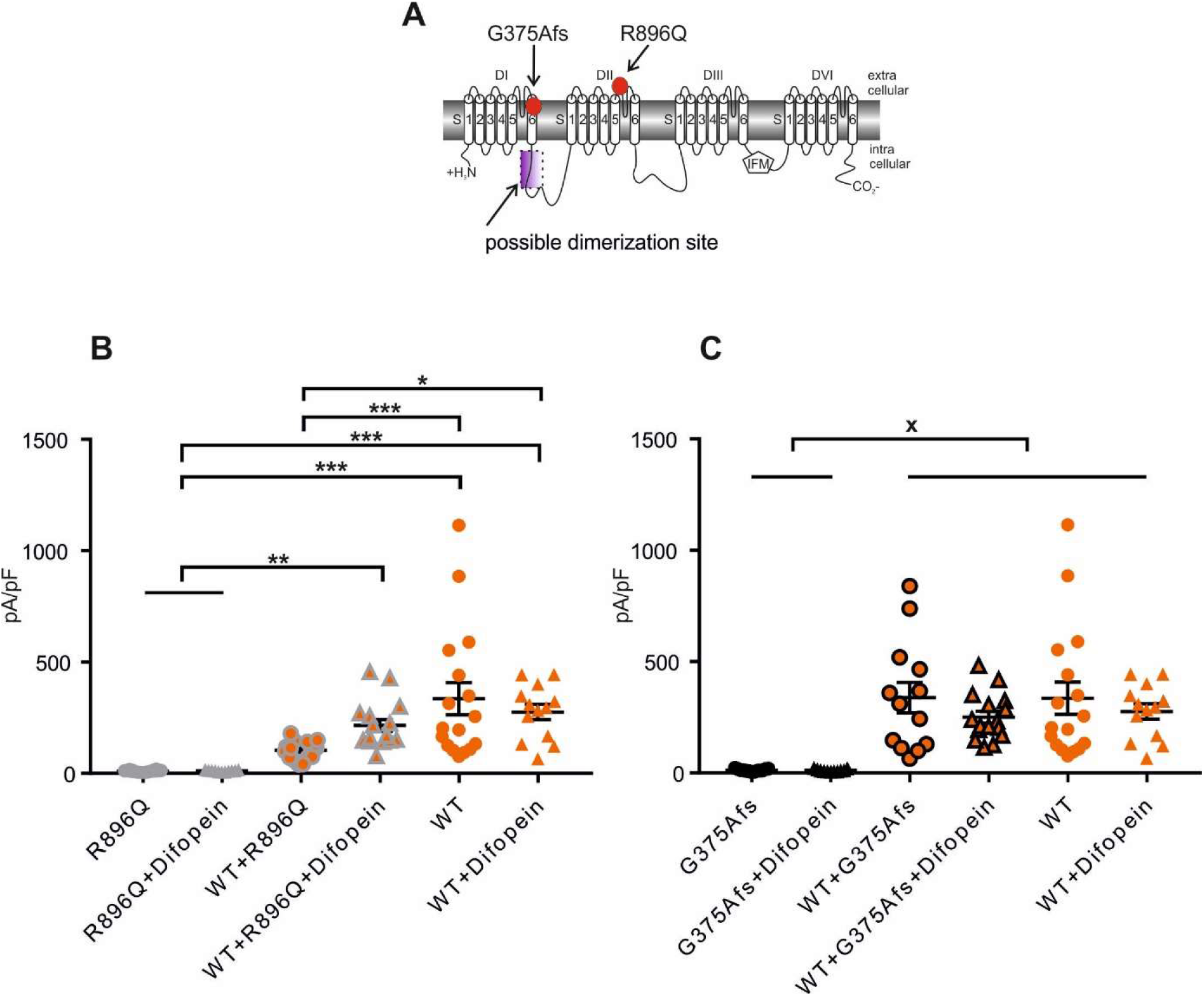
Loss-of-function mutations confirm channel dimerization of hNav1.7. A) Schematic representation hNav1.7 with the localization of hNav1.7/G375Afs, hNav1.7/R896Q (red dots) and the possible dimerization site in the linker region between DI and DII (purple rectangle). B) Current density in pA/pF: hNav1.7/R896Q =10.78±1.08, N=16; hNav1.7/R896Q+difopein: 11.93±1.01, N=11; WT+hNav1.7/R896Q: 104.2±10.25, N=16; WT+hNav1.7/R896Q+difopein: 216.2±26.27, N=16; WT=335.6±72.28, N=17; WT+difopein: 276.2±34.11, N=13 *p<0.05, **p<0.01, ***p<0.001. For exact p-values, please refer to supplementary table 1. C) Current density in pA/pF: hNav1.7/G375Afs: 12.36±1.62,N=13; hNav1.7/G375Afs+difopein: 11.27±1.69, N=11; WT+hNav1.7/G375Afs: 338±68.65, N=13, WT+hNav1.7/G375Afs+difopein: 250.7±26.01, N=16; WT: 335.6±72.28, N=17; WT+difopein: 276.2±34.11, N=13; x indicates significant changes. For exact p-values, please refer to supplementary table 2. Data are shown as mean ± SEM.

First, we confirmed the previously reported absence of sodium current in the missense mutation of the full length hNav1.7/R896Q (Fig. 4B, Cox et al. 2010). Co-expression of hNav1.7/R896Q in the WT cell line led to a significantly reduced current density (Fig. 4B). Adding difopein, which is supposed to interfere with functional channel coupling, partially restored the current density, suggesting that a dimerization between hNav1.7/R896Q and WT can be responsible for the reduction in current density. Difopein has no impact on the current density of hNav1.7 WT (Fig. 4B).

The truncation mutation hNav1.7/G375Afs abolishes sodium current (Fig. 4C), in line with the symptoms of the patient (Shorer *et al*, 2014). Interestingly, co-expression of hNav1.7/G375Afs in the Nav1.7 WT cell line did not affect the current density neither with nor without difopein (Fig. 4C). Thus, it is likely that the truncated channels of the hNav1.7/G375Afs are not able to interact with the full length WT Nav1.7 channel to form dimers (see discussion), suggesting that the potential dimerization site resides between the two mutations in the DI-DII linker, a similar region as proposed for hNav1.5 before (Clatot et al. 2017).

Now that we have established evidence for functional dimerization of hNav1.7 we set out to investigate how this may affect the allosteric inactivation mechanism: We co-expressed difopein with hNav1.7/A1632E in order to suppress functional dimerization. The presence of difopein cut the persistent current of hNav1.7/A1632E in half, reducing it from 14.9 ± 1.4% without to 7.0% ± 0.6% with difopein (Fig. 5 A+B). Thus, functional dimers of Nav1.7 seem to affect the disease causing persistent current of hNav1.7/A1632E, adding additional pathophysiological evidence to channel dimerization.

**Figure 5:**
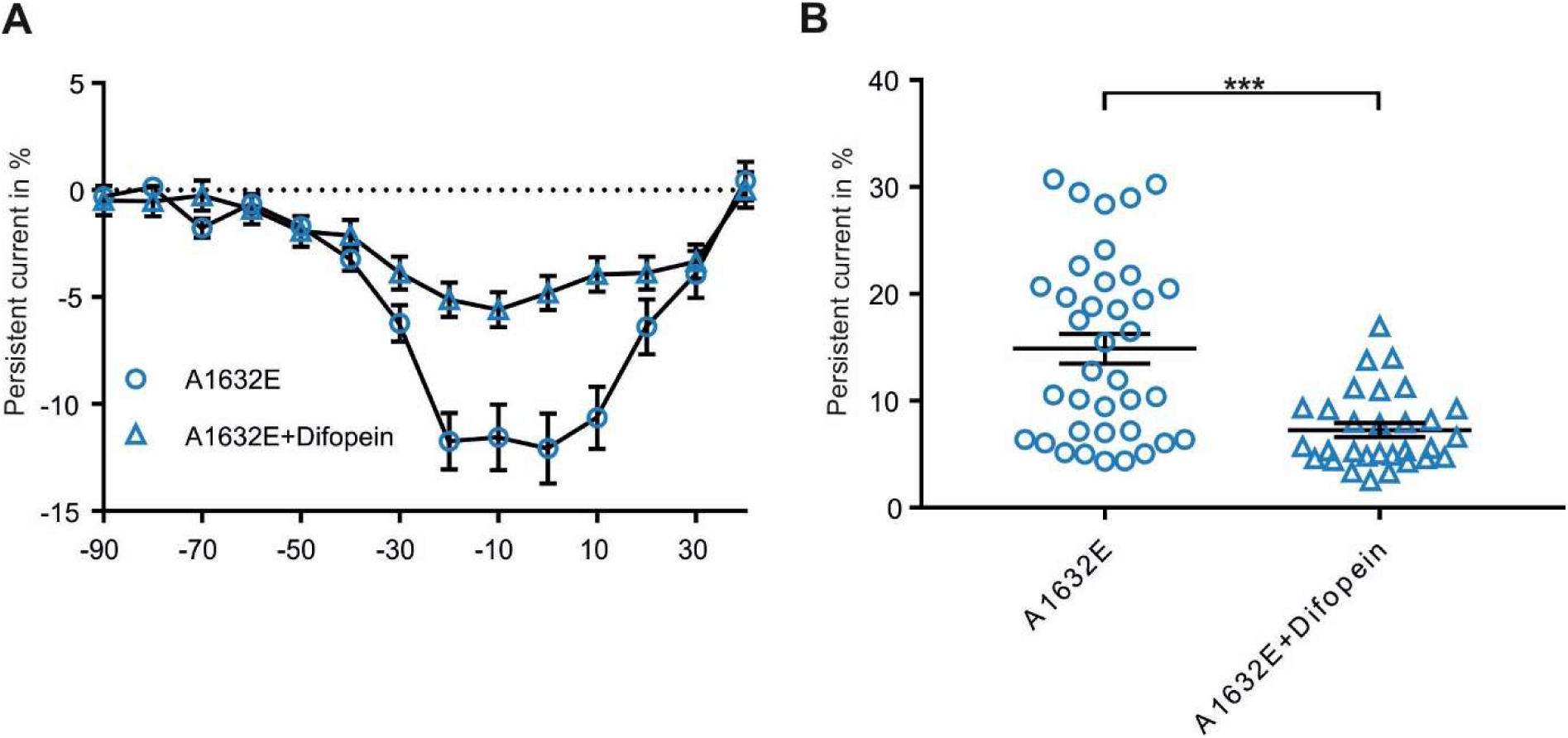
Difopein reduces the persistent current of hNav1.7/A1632E. A) Relative persistent current in percent of peak current for each test-pulse voltage. The measured persistent current at each voltage step was normalized towards the peak inward current of the cell. B) Maximal persistent current for Nav1.7/A1632E: 14.87%±1.39%, N=37; hNav1.7/A1632E+difopein: 7.03%±0.63%, N=29. ***p<0,0001. Data are shown as mean ± SEM.

### Dimerization affects physiological gating of hNav1.7

Our results suggest that channel dimerization affects fast inactivation of the mutant channel hNav1.7/A1632E. To assess a possible physiological relevance of VGSC dimerization in humans under physiological conditions, we assessed the steady-state fast inactivation of WT hNav1.7. The presence of difopein in the WT Nav1.7 cell line introduced a 2.9 mV shift of V_1/2_ towards more hyperpolarized potentials (Fig. 6A+B), suggesting that WT hNav1.7 not only dimerizes but might actually gate as a functionally coupled dimer. Interference with 14-3-3 binding via difopein allows for regulation of steady-state fast inactivation under physiological conditions.

**Figure 6:**
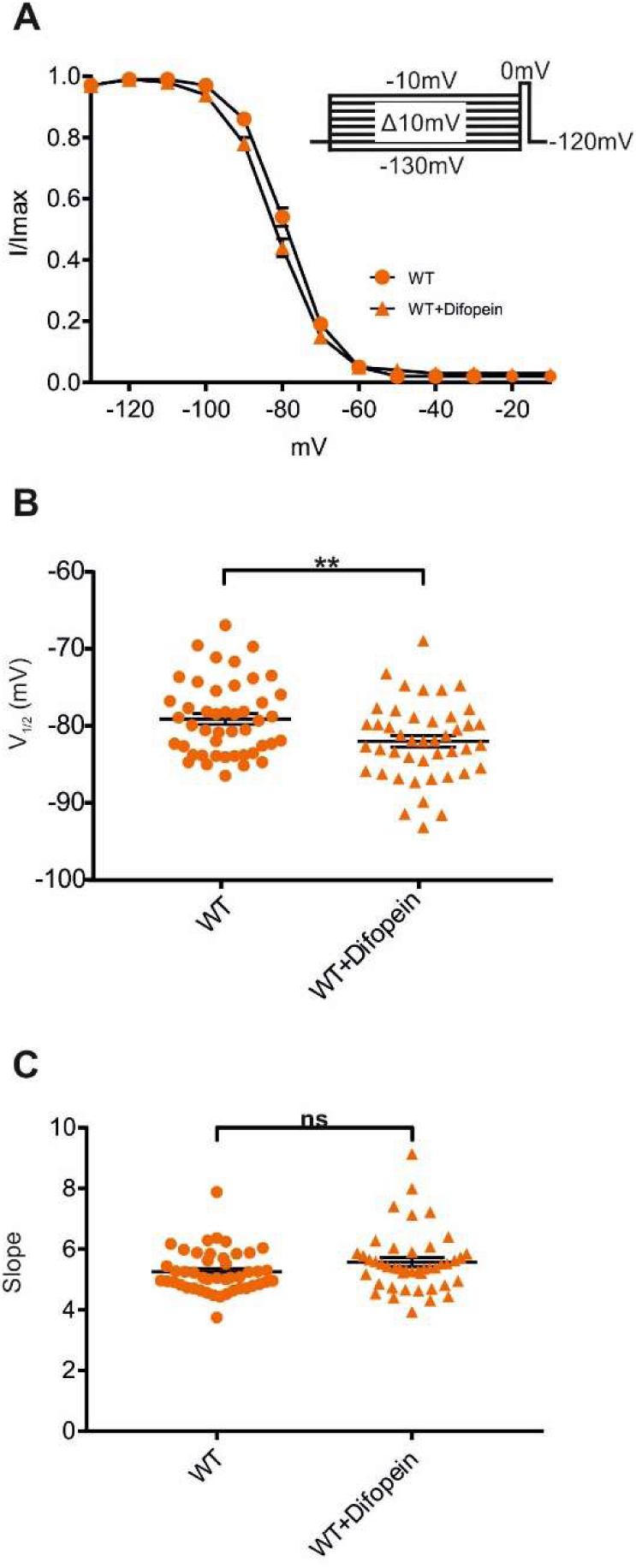
Dimerization affects hNav1.7 fast inactivation. A) Voltage dependence of fast inactivation of Nav1.7/WT ± difopein. Voltage-clamp protocol is depicted as inset. B) Half-maximal (V_1/2_) voltage dependence of fast inactivation: WT: −79.12mV±0.71, N=46; WT+difopein: −82mV±0.76, N=44. **p=0,0067. C) Slope of fast inactivation: WT: 5.25±0.1, N=46; WT+difopein: 5.57±0.15, N=44. p=0.1. Data are shown as mean ± SEM.

## Discussion

In the present study we show that the pain-linked mutation hNav1.7/A1632E impairs the allosteric fast inactivation mechanism of the channel, we demonstrate that Nav1.7 forms functional dimers and that uncoupling dimers reduces disease-relevant persistent current of the hNav1.7/A1632E mutation.

### Allosteric fast inactivation is impaired by hNav1.7/A1632E leading to persistent currents

The prominent persistent current of hNav1.7/A1632E is likely to underlie the symptoms associated with PEPD (Eberhardt *et al*, 2014). In general, persistent current is thought to occur due to impaired fast inactivation (Dib-Hajj *et al*, 2013). Until very recently, fast inactivation was believed to result from the IFM-motif directly blocking the channel pore (Goldin, 2003; West *et al*, 1992), but the recently published cryo-EM channel structures suggest an allosteric mechanism for IFM binding-induced fast inactivation (Pan *et al*, 2018; Shen *et al*, 2017; Yan *et al*, 2017; Shen *et al*, 2019; Clairfeuille *et al*, 2019; Xu *et al*, 2019). According to this allosteric mechanism, the IFM motif binds to a hydrophobic pocket which is built by residues including A1632 on the S4–S5 linker (Fig. 2A). Substitution of A1632 with glutamate adds additional size, which may sterically impair IFM binding, but it also increases hydration of the pocket. This seems to severely interfere with IFM binding and thus likely causes the prominent persistent current of hNav1.7/A1632E.

Interestingly, replacing A1632 with aspartate, which is smaller than glutamate but also carries a negative charge, induces even larger persistent currents (Eberhardt *et al*, 2014). Substitution with the uncharged amino acids threonine (Eberhardt *et al*, 2014) or glycine (Yang *et al*, 2016) left the persistent current unaltered, but shifted voltage dependence of steady-state fast inactivation to more depolarized potentials. In contrast, conservative side chain substitutions such as A1632V had no effect on either persistent current or steady-state fast inactivation (Eberhardt *et al*, 2014). Thus, it seems that introduction of negative charge in the IFM binding site interferes with fast inactivation resulting in persistent currents and our findings support the notion that disruption of the hydrophobic binding pocket of the IFM motif impairs fast inactivation.

### Nav1.7 forms functional dimers

Our data show that next to hNav1.5 and hNav1.2, hNav1.7 also forms functional dimers. Two binding sites for 14-3-3 were reported: One between amino acids 416 - 467, the second one between amino acids 517 – 555, both on the DI-DII linker (Supplementary Figure S2). In addition, a 14-3-3 independent channel interaction site, the so-called α-α subunit interaction site, was reported to reside between these two 14-3-3 binding sites (Clatot *et al*, 2017). In fact, we suggest that channel-dimerization may be considered as a general feature of VGSC, since alignment of the suggested dimerization sites revealed conserved amino acids for almost all subtypes in comparison to hNav1.5 except for hNav1.4. (Supplementary Figure S2).

Fast inactivation of hNav1.7 WT is modified by difopein, suggesting that regulation by interaction with 14-3-3 may function as a modulator of channel gating under physiological conditions. Depolarizing voltage-depencence of fast inacitvation to a similar extent as that observed by difopein expression with Nav1.7 WT were reported before to be induced by mutations identified in pain patients (e.g. Wu *et al*, 2013), revealing that very small changes can have a significant impact on cellular excitability. The observed shift for WT fast inactivation could hence lead to an altered physiological pain perception influenced by VGSC dimerization-pattern. Modulating availability and binding of 14-3-3 could change these dimerization-patterns, possibly due to varying phosphorylation, heterodimer formation between different 14-3-3 isoforms or post-translational modifications (Pennington *et al*, 2018). Expression of 14-3-3 was shown in DRG growth cones, suggesting that 14-3-3 may play a regulatory role in VGSC dimerization within nociceptive nerve endings (Kent *et al*, 2010).

### Dimerization affects severity of pain-linked mutation induced hNav1.7 gating changes

Inhibiting dimerization drastically reduced the persistent current of hNav1.7/A1632E (Fig. 5), showing that dimerization can regulate the severity of mutation induced gating changes. Since gain-of-function mutations lead to symptoms in heterozygous carriers, (Eberhardt *et al*, 2014; Lampert *et al*, 2010; Emery *et al*, 2015; Yang *et al*, 2016; Klein *et al*, 2013; Estacion *et al*, 2008; Meents *et al*, 2019) new questions arise on how dimerization patterns of homo- and/or heterodimers influence disease manifestation and progress, not only of the here reported pain-linked mutation, but of many more known to date.

In loss-of-function mutations however, heterozygous conditions remain without obvious symptoms (Cox *et al*, 2010; Goldberg *et al*, 2007; Klein *et al*, 2013; Cox *et al*, 2006; Emery *et al*, 2015; McDermott *et al*, 2019). In the case of hNav1.7/R896Q the absence of sodium current is explained by deficient channel trafficking (Cox *et al*, 2010). Although some of the mutant channels reach the cell membrane, the percentage does not suffice for the occurrence of sodium current. Interestingly, for hNav1.5, it was shown that the co-expression of a trafficking deficient mutation with WT reduced channel expression on the cell membrane (Wang *et al*, 2015). There might be parallels between these findings and those reported here on the hNav1.7/R896Q mutation. Co-expression of the mutation and dimerization with the WT could retain the functioning channels within the cell. This would explain why sodium current increases in the presence of difopein although the transfected cDNA-concentration remains stable. The absence of dimerization could allow the WT channel to travel to the cell membrane, causing more sodium current to occur. We showed that although the current density was significantly reduced in the presence of WT and hNav1.7/R896Q, a small sodium current was still present. Our results go along with a recent study on hNav1.7 in iPSC-derived nociceptors of CIP patients: The authors showed that in neurons with a bi-allelic expression of a CIP mutation, restoring one deficient hNav1.7 allele was enough to regain some but not all of the electrophysiological functions of these neurons (McDermott *et al*, 2019). Channel dimerization affects the functioning channels in a dominant-negative way but the remaining channel function seems still sufficient for pain perception. Thus, it appears that one Nav1.7 allele produces enough functioning channels to generate sodium current and support pain perception, and therefore CIP mutations occure in patients always as homozygous or compound heterozygous mutations.

The truncation mutation hNav1.7/G375Afs, on the other hand, does not seem to dimerize with the WT likely because the mutant channel is truncated before of the assumed dimerization site (Clatot *et al*, 2017). It is also possible that the mutant channel did not express adequately, and that dimerization was thus absent. However, since our other mutants expressed sufficiently and 14-3-3 is present in our expression system (Supplementary Figure S1), we argue that the conditions for dimerization were given and that the abolishment of the putative dimerization site is likely to be the reason for missing effects of difopein.

Pain syndromes are complex diseases: different phenotypes appear in one family bearing the same mutation (Michiels *et al*, 2005) and the exact relation between electrophysiology and phenotype still remains uncertain (Hampl *et al*, 2016; Emery *et al*, 2015). Channel-dimerization is a new aspect to be considered when explaining pain syndromes and their complex phenotypes. Additionally, even diseases caused by different Nav-subtypes might need to be re-evaluated, considering the possibility of channel dimerization.

### Conclusion

Here, we show that hNav1.7 forms functionally relevant dimers, that this dimerization modifies the mutation-induced phenotype of the pain-linked hNav1.7/A1632E substitution, and that this modification most likely appears via the allosteric inactivation mechanism suggested by recently published VGSC structures. Our work supports the concept of sodium channel dimerization, its physiological function and reveals its relevance to human pain syndromes.

## Materials and Methods

### Plasmids

A hNav1.7 plasmid in a pCMV6-neo vector was used as previously reported (Klugbauer *et al*, 1995; Stadler *et al*, 2015). hNav1.7/R896Q and hNav1.7/G375Afs were both introduced into a pCMV6-neo vector. hNav1.7/A1632E was used in a pCDNA3 vector as previously reported (Eberhardt *et al*, 2014). Difopein in a pIRES2-EGFP Vector (BioCat GmbH, Heidelberg, Germany) and GFP in a pMax vector (Lonza, Basel, Switzerland) were purchased.

Plasmids for cRNA-encoded protein expression in oocytes were designed with the Vector NTi Deluxe v4.0 software (InforMax). The full-length coding region of hNav1.5 and hNav1.7 were individually subcloned into the oocyte expression vector pNKS2 (Gloor *et al*, 1995) by Gateway PCR and In-Fusion HD Cloning (Takara Bio Europe, Göteborg, Sweden), respectively. For oocyte expression, the difopein coding region was subcloned from the pIRES2-EGFP vector into the pNKS2 vector by megaprimer PCR. Human transient receptor potential cation channel subfamily V member 1 (hTrpV1, transcript variant 3) in pDONR201 (DNASU plasmid ID HsCD00081472 corresponding to NCBI reference sequence NM_080706.3) was purchased from the DNASU Plasmid Repository (Tempe, Arizona, USA), subcloned into the pNKS2 vector by Gateway cloning. hNav1.5, hNav1.7 and hTRPV1 were fused in frame to a C-terminal GFP coding region using megaprimer PCR (Kirsch & Joly, 1998; Perez *et al*, 2006) to yield hNav1.5^GFP^, hNav1.7^GFP^ and hTRPV1^GFP^. Cloning and propagation of hNav1.7, but not the other plasmids, required the use of stable competent E. coli (NEB, Frankfurt, Germany) suited for plasmids with instable inserts. All constructs were verified by restriction patterns and commercial DNA sequencing (MGW, Ebersberg, Germany).

### Cell Culture and Transfection

Two human embryonic kidney cell (HEK cell) lines were used: HEK293T cells and a HEK cell line stably expressing WT hNav1.7 (Hampl *et al*, 2016; Körner *et al*, 2018). To the second cell line we refer to as WT cell line. Cells were cultured in Dulbecco’s modified Eagle’s medium (DMEM; Gibco–Life Technologies, Carlsbad, CA, USA) containing 1.0 g l−1 glucose and 10% fetal bovine serum (Gibco–Life Technologies, Carlsbad, CA, USA). Medium for the WT cell line contained 1% Geneticin G418 (A&E Scientific, Marcq, Belgium) as a selection marker. Cells were incubated at 37°C and 5% C0_2_. Transfection was performed using Jet-PEI reagent (POLYPLUS Transfection, Illkirch, France). For all conditions, a total amount of 1.5 µg DNA was used.

For single transfections of the hNav1.7 mutants, 1.25 µg of plasmid were transfected into HEK293T cells. Either 0.25 µg GFP or Difopein were added. hNav1.7/R896Q and hNav1.7/G375Afs were compared to the WT cell line transfected with either 1.5 µg GFP or difopein. For introduction into the WT cell line, 1.25 µg hNav1.7/R896Q or hNav1.7/G375Afs were used. 0.25 µg GFP or difopein were added. For experiments concerning the steady-state fast inactivation of the WT, the WT cell line ± 1.5 µg difopein was used.

### Electrophysiology

Only green fluorescent cells were used for whole-cell patch clamp recordings 27-60 hours after transfection using a HEKA EPC 10 USB patch-clamp amplifier (HEKA Electronics, Lambrecht, Germany). Sampling rate was set to 10kHz. Patch Pipettes were pulled with a DMZ pipette puller (ZEITZ Instrumente Vertrieb GmbH, Martinsried, Germany) and pipettes with a tip resistance in-between 0.8 and 3.0 MΩ were used. All recordings were performed at room temperature (21°C±2C) and the liquid junction potential was not corrected.

For patch clamp recordings, the following bath solution was used: 140mM NaCl, 3mM KCl, 1mM MgCl_2_, 1mM CaCl_2_, 10mM HEPES, 20mM Glucose. PH was adjusted to 7.4 using NaOH, the osmolarity of the solution was 310±10 mOsm. Internal pipette solution contained the following: 10mM NaCl,140mM CsF, EGTA 1mM, 10mM HEPES, 18mM Sucrose. The pH was adjusted to 7.33 and the osmolarity was 310±10 mOsm.

For HEK cell recordings, series resistance was below 7MΩ at any time for all of the cells and compensated to ≥65%. Leak current subtraction was performed online via a P/4 procedure. After reaching whole cell configuration, cells were held at a holding potential of −120mV for 3min and pulsed with 0.1Hz to allow stabilization of the inward current. Immediately afterwards, the current-voltage-relationship was measured by stepwise 40ms depolarizations from −90mV to +40mV in 10mV increments.

A Boltzmann equation was used for fitting: G/G_max_=(G_max_-G_min_)/(1+exp[(V_1/2_−V_m_)/k]). (G_max_= maximum sodium conductance, V_1/2_= membrane potential at half maximal activation, V_m_= Membrane potential, k= slope factor). G_max_ was set to 1 and G_min_ to 0.

Fast inactivation was measured 10 s after the end of the activation protocol by applying a test-pulse to 0mV for 40ms after pre-pulses of 500ms. Pre-pulses were increased in 10mV increments from - 130mV to −10mV. Cells were held at a holding potential of −120mV. The same equation as shown above was used for fitting. I_max_ was set to 1, I_min_ was not defined.

The persistent current measured between 34ms and 39.6ms of each 40ms test-pulse was normalized to the transient peak inward current of the same cell. The maximum persistent current of each cell was then used for comparison.

### Polymerase Chain Reaction (PCR)

PCR was performed with RNA extracted from untransfected HEK293T cells and the WT cell line. For extraction, a Nucleospin RNA kit from Macherey-Nagel (Düren, Germany) was used and the instructions were followed as provided by the company. cDNA was synthesized with a sensifast cDNA synthesis kit by bioline (London, UK). To test for different 14-3-3 isoforms, seven different human-primers fabricated by eurofins genomics (Ebersberg, Germany) were used (Table1). Taq-Polymerase, Thermopol Buffer and dNTP from New England BioLabs inc. (Frankfurt am Main, Germany) were used. The protocol included 35 cycles and an annealing temperature at 52°C.

**Table 1:**
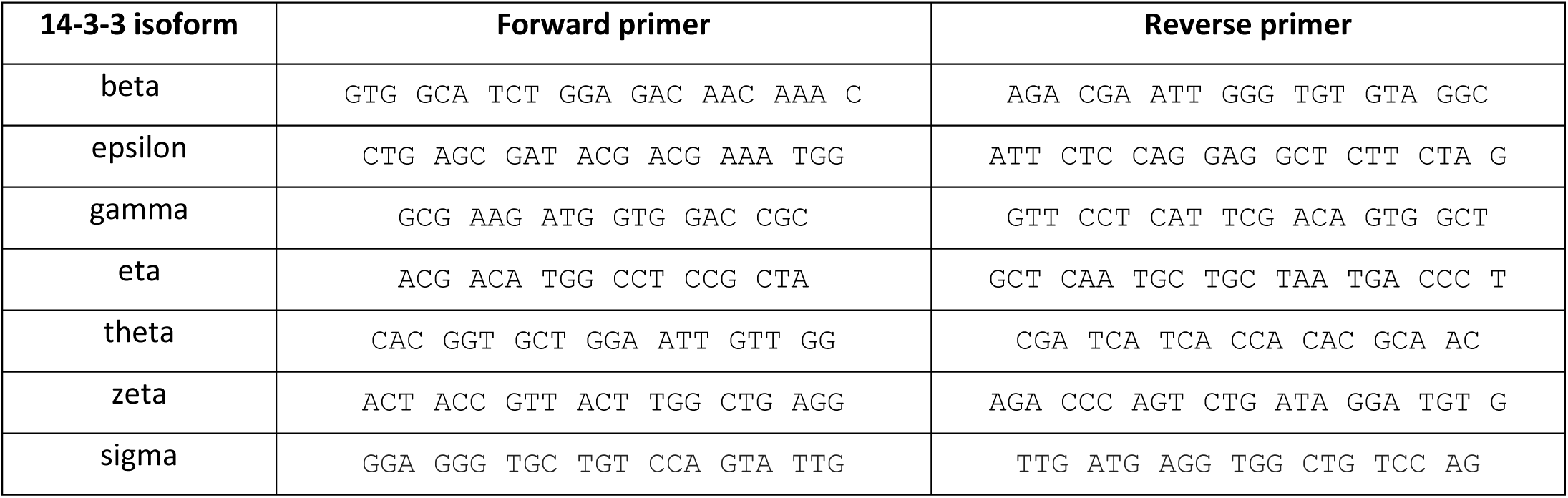
Forward and reverse primers used in the PCR are given for each 14-3-3 isoform.

### Biochemical analysis of hNav1.5^GFP^ and hNav1.7^GFP^ expressed in X. laevis oocytes

For a biochemical characterization of hNav1.7, collagenase-defolliculated *X. laevis* oocytes (Dumont stages V and VI) were used as previously described (Stolz *et al*, 2015). The oocytes were injected with capped and polyadenylated cRNA (30-60 ng/oocyte) and then maintained at 19 °C in oocyte Ringer’ solution (ORi) consisting of 90 mM NaCl, 1 mM KCl, 1 mM CaCl_2_, 1 mM MgCl_2_ and 10 mM Hepes/NaOH pH 7.4 supplemented with 50 µg/ml of gentamycin. At day 2 post-cRNA injection, the oocytes were homogenized in ice-cold 0.1 M sodium phosphate, pH 8.0 (10 µl/oocyte) supplemented with 1 mM Tris(2-carboxyethyl)phosphine HCl (TCEP) and protease inhibitors (pepstatin, leupeptin, antipain, Pefablock SC) plus one of the following non-ionic detergents: digitonin (water soluble quality, Serva, Heidelberg, Germany), n-dodecyl ß-D-maltoside (DDM, AppliChem, Darmstadt, Germany), glyco-diosgenin or lauryl maltose neopentyl glycol (GDN or NG310, respectively; Anatrace, Maumee, USA) in concentrations as indicated in the figures. After two clearing spins at 10.000 g for 15 min each, aliquots of the supernatant were resolved by hrCN-PAGE electrophoresis (Wittig *et al*, 2006) and clear SDS-urea-PAGE (Stolz *et al*, 2015; Fallah *et al*, 2011). The gels were destained by overnight incubation in acetonitrile/ammonium carbonate, washed twice in 0.1 M sodium phosphate pH 8.0 (Wolf *et al*, 2011), and the wet PAGE gels were scanned on a Typhoon fluorescence scanner (GE Healthcare). Figures were prepared with halftone images by using the Image-Quant TL software v8.2 (GE Healthcare) for contrast adjustments, Adobe Photoshop CS 8.0 for level adjustment and cropping, and Microsoft PowerPoint 2000 for labeling.

### Molecular dynamics simulations

All-atom MD simulations of the human Nav1.7 alpha/beta1 subunit complex (PDB ID: 6J8I) (Shen *et al*, 2019) were carried out using the CHARMM36m force field with the TIP3P water model (Huang *et al*, 2017). The simulation box contained an equilibrated 1-palmitoyl-2-oleoyl-sn-glycero-3-phosphocholine (POPC) lipid bilayer, surrounded by 250 mM NaCl aqueous solution. The DIII–DIV linker containing the IFM motif was removed from the channel structure, and Nav1.7 was modelled using the residue ranges 114–417/715–961/1164–1757 for the alpha subunit and 20–192 for the β1 subunit (using isoform 3 residue numbering). The complex was embedded into the bilayer using *g_membed* (Wolf *et al*, 2010). Glu180 and Asp1256 were assigned a protonated state, all other amino acids were modelled in their default ionization state to reflect the most probable state at neutral pH based on pKa calculations using PROPKA 3.1 (Søndergaard *et al*, 2011; Olsson *et al*, 2011).

We performed MD simulations using GROMACS 2018 (Abraham *et al*, 2015) in the NPT ensemble with periodic boundary conditions and an integration time step of 2 fs. Temperature was maintained at 310 K using the velocity-rescaling thermostat; pressure was kept at 1 bar using the semi-isotropic Parrinello-Rahman barostat as described recently (Machtens *et al*, 2015). Lennard-Jones interactions were truncated at 12 Å with a force switch smoothing function from 10–12 Å. Electrostatic interactions were calculated using particle mesh Ewald method and a real space cutoff of 12 Å. The simulation systems of wild-type and A1632E Nav1.7 were equilibrated with position restraints on the protein heavy atoms for 500ns, followed by ∼20ns with backbone-only position restraints, and 200-ns production runs without any restraints. We used MODELLER to insert the A1632E side chain substitution (Webb & Sali, 2016).

Water densities were calculated from the production runs in the time window 10–200 ns using GROmaps (Briones *et al*, 2019)and visualized by mapping water densities onto the water-accessible protein surface of the IFM-binding pocket. PyMOL was used to render all molecular images (PyMOL).

### Sequence alignment

The putative dimerization sites of hNav1.5 (Clatot *et al*, 2017) were aligned with different human VGSC subtypes using UniProt and BLAST (Altschul *et al*, 1990; Bateman, 2019).

## Data analysis

For data analysis, Fitmaster software (HEKA electronics, Lambrecht, Germany) and IgorPro software (WaveMetrics, Portland OR, USA) were used. Statistical analysis was performed with Prism 7 (GraphPad, San Diego CA, USA). Groups were tested for a Gaussian distribution using a D’Agostino Pearson test. In case of non-parametric testing, a Mann-Whitney test was applied; in case of parametric testing an unpaired t-test was used. When comparing more than two groups, an ordinary one-way ANOVA with Bonferroni post-hoc correction was used. To quantify variation, the standard error of the mean (SEM) is displayed. Concerning the persistent current and current density, outliers were checked for significance using Grubb’s test (GraphPad QuickCals, San Diego CA, USA) and excluded from analysis. Data are shown as mean ± SEM for all figures.

## Acknowledgments

We thank Brigitte Hoch for excellent technical support.

This work was supported by the Deutsche Forschungsgemeinschaft (DFG, German Research Foundation) to J.P.M (MA 7525/1-1; as part of the Research Unit FOR 2518, DynIon; project P4) and to A.L. (LA 2740/3-1, RTG 2416 MultiSenses-MultiScales and the RTG 2415 Mechanobiology of 3D epithelial tissues (ME3T)). The authors gratefully acknowledge the computing time granted by the JARA-HPC Vergabegremium and VSR Commission on the supercomputer JURECA (Krause & Thörnig, 2016) at Forschungszentrum Jülich.

## Author contributions

All authors participated in writing and have critically revised and approved the final version of the manuscript.

A.R. participated in study design, performed and analyzed PCR, patch-clamp experiments, discussed and interpreted the results of the study and wrote the manuscript.

J.K. performed protein structural analysis, discussed and interpreted the results of the study.

N.B. performed the biochemical experiments.

S.D. subcloned the vectors for oocyte expression and performed biochemical experiments.

P.H. performed mutagenesis and performed and analyzed the PCR.

C.A.B. performed and analyzed molecular dynamics simulations.

J.M. designed and interpreted the patch-clamp experiments and discussed and interpreted the results of the study.

J.P.M. advised on the setup and analysis of molecular dynamics simulations, and discussed and interpreted the results of the study.

G.S. designed the biochemical experiments, analyzed and interpreted the results, and discussed and interpreted the results of the study.

A.L. conceived and designed the study, discussed and interpreted the results of the study.

All authors agree to be accountable for all aspects of the work in ensuring that questions related to the accuracy or integrity of any part of the work are appropriately investigated and resolved. All persons designated as authors qualify for authorship and all those who qualify for authorship are listed.

## Supplementary Information

**Supplementary Figure S1:**
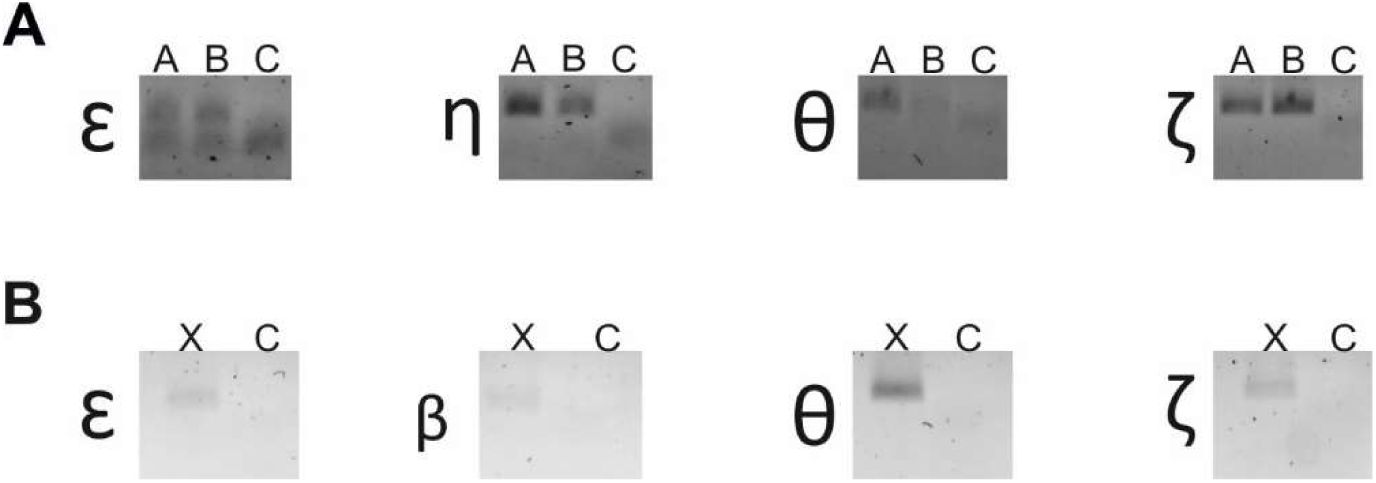
Expression of 14-3-3 isoforms in HEK cells and X.leavis oocytes. A) PCR of 14-3-3 isoform mRNA expression in WT cell line (lane A) and untransfected HEK293T cells (lane B) in comparison to water control (lane C). B) PCR of 14-3-3 isoform mRNA expression in X.laevis (lane X) in comparison to water control (lane C).

**Supplementary Figure S2:**
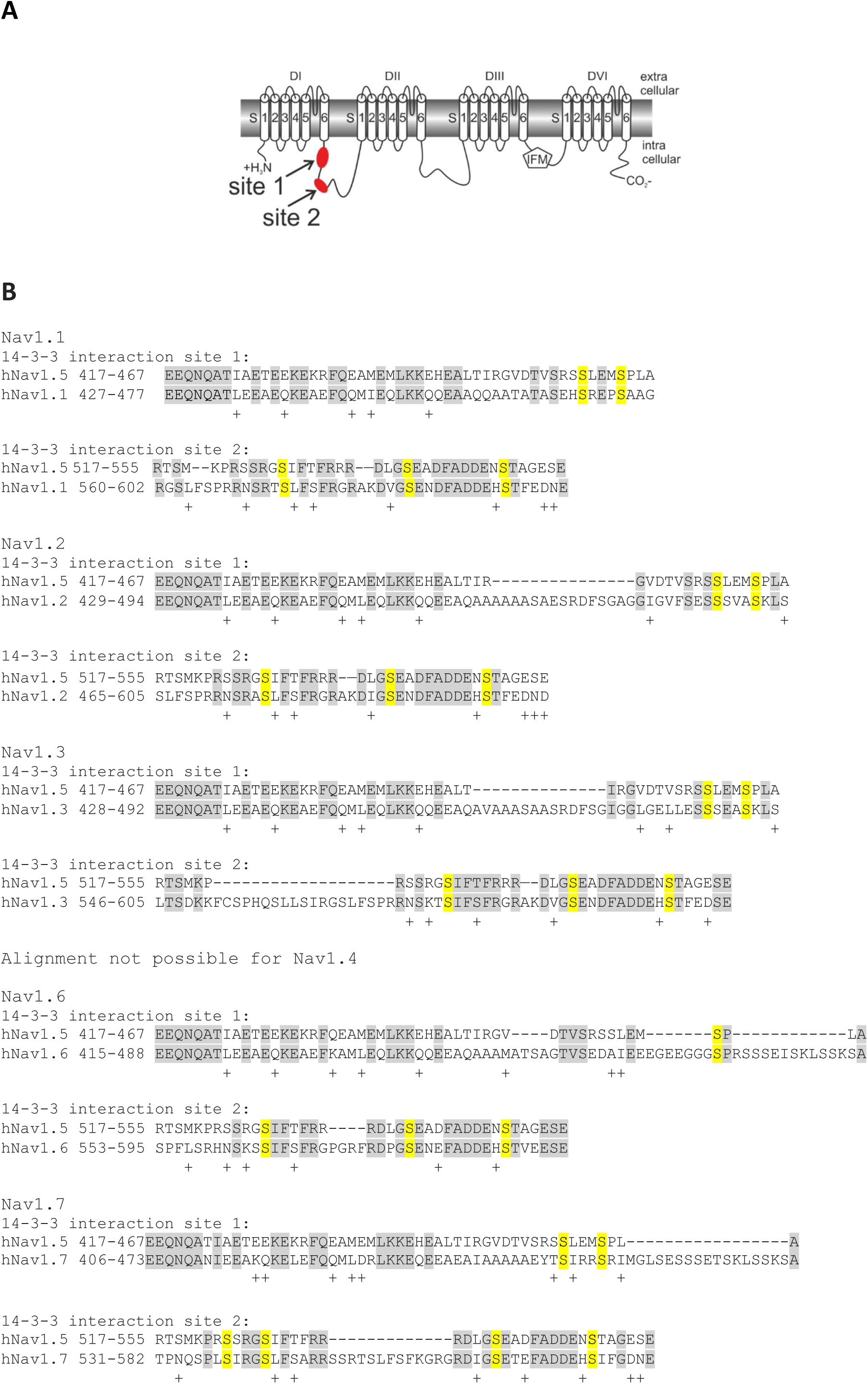
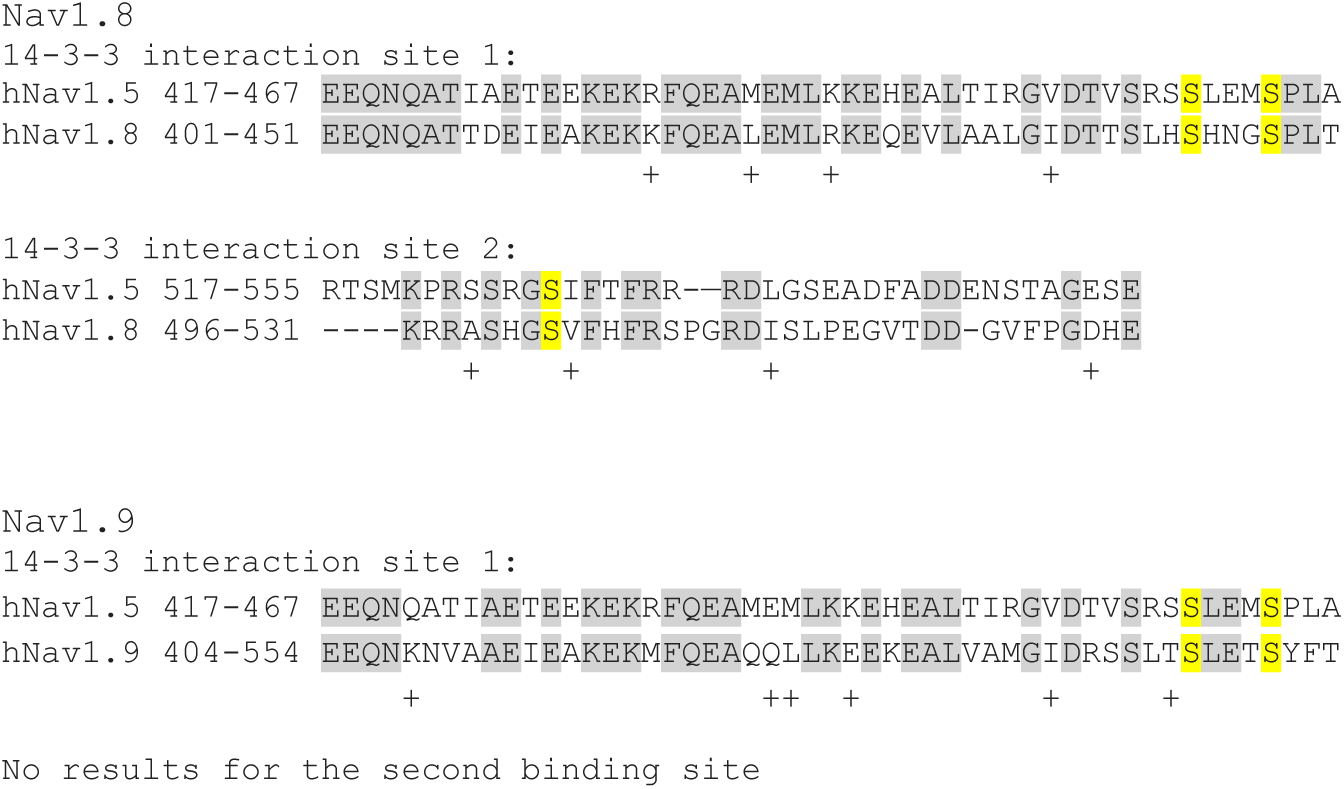
Alignment of dimerization site of the hNav1.5 and hNav1.7. A) Schematic view of the suggested 14-3-3 interaction sites that support dimerization. B) The sequence of the end of the S6 segment of domain I and the beginning of the linker between domain I – domain II is shown for Nav1.5 and all known Navs. Conserved amino acids are shaded in grey and similar amino acids are marked by a “+”. Conserved serines for Nav1.5 and Nav1.7 are marked in yellow as those residues may act as interaction sites for 14-3-3. Alignment was made using UniProt (McGARVEY *et al*, 2008) and BLAST (Altschul *et al*, 1990).

**Supplementary Table 1:** p-values for the different conditions using hNav1.7/R896Q and WT

| Compared groups | p-value |
| --- | --- |
| WT+R896Q vs. WT+Difopein | *p=0.0358 |
| R896Q vs. WT+ R896Q+Difopein | **p=0.0024 |
| R896Q+Difopein vs. WT+R896Q+Difopein | **p=0.0097 |
| WT+R896Q vs. WT | ***p=0.0003 |
| R896Q vs. WT | ***p<0.0001 |
| R896Q+Difopein vs. WT | ***p<0.0001 |
| R896Q vs. WT+Difopein | ***p<0.0001 |
| R896Q+Difopein vs. WT+Difopein | ***p=0.0005 |

**Supplementary Table 2:** p-values for the different conditions using hNav1.7/G375Afs and WT

| Compared groups | p-value |
| --- | --- |
| G375Afs vs. WT+G375Afs+Difopein | *p=0.0102 |
| G375Afs+Difopein vs. WT+G375Afs+Difopein | *p=0.0165 |
| G375Afs vs. WT+Difopein | **p=0.0054 |
| G375Afs+Difopein vs. WT+Difopein | **p=0.0087 |
| G375Afs vs. WT | ***p<0.0001 |
| G375Afs vs. WT+G375Afs | ***p=0.0002 |
| G375Afs+Difopein vs. WT | ***p=0.0002 |
| G375Afs+Difopein vs. WT+G375Afs | ***p=0.0005 |

